# 3D-Reconstruction of the Human Conventional Outflow System by Ribbon Scanning Confocal Microscopy

**DOI:** 10.1101/2020.01.17.910257

**Authors:** Ralitsa T. Loewen, Susannah Waxman, Chao Wang, Sarah Atta, Si Chen, Simon C. Watkins, Alan M. Watson, Nils A. Loewen

## Abstract

**Purpose:** Risk for glaucoma is driven by the microanatomy and function of the anterior segment. We performed a computation-intense, high-resolution, full-thickness ribbon-scanning confocal microscopy (RSCM) of the outflow tract of two human eyes. We hypothesized this would reveal important species differences when compared to existing data of porcine eyes, an animal that does not spontaneously develop glaucoma.

**Methods:** After perfusing two human octogenarian eyes with lectin-fluorophore conjugate and optical clearance with benzyl alcohol benzyl benzoate (BABB), anterior segments were scanned by RSCM and reconstructed in 3D for whole-specimen rendering. Morphometric analyses of the outflow tract were performed for the trabecular meshwork (TM), limbal, and perilimbal outflow structures and compared to existing porcine data.

**Results:** RSCM provided high-resolution data for IMARIS-based surface reconstruction of outflow tract structures in 3D. Different from porcine eyes with an abundance of highly interconnected, narrow, and short collector channels (CCs), human eyes demonstrated fewer CCs which had a 1.5x greater cross-sectional area (CSA) and 2.6x greater length. Proximal CC openings at the level of Schlemm’s canal (SC) had a 1.3x larger CSA than distal openings into the scleral vascular plexus (SVP). CCs were 10.2x smaller in volume than the receiving SVP vessels. Axenfeld loops, projections of the long ciliary nerve, were also visualized.

**Conclusion:** In this high-resolution, volumetric RSCM analysis, human eyes had far fewer outflow tract vessels than porcine eyes. Human CCs spanned several clock-hours and were larger than in porcine eyes. These species differences may point to factors downstream of the TM that increase our vulnerability to glaucoma.

**Grant information:** National Eye Institute K08EY022737 (NAL); Initiative to Cure Glaucoma of the Eye and Ear Foundation of Pittsburgh (NAL); Wiegand Fellowship of the Eye and Ear Foundation of Pittsburgh (YD); P30-EY08098 (NAL); Department grant by Research to Prevent Blindness (NAL); an unrestricted fellowship grant from the Xiangya Hospital of Central South University (SC).

## Introduction

Recent experiments in ex vivo human [1] and pig [1, 2] eyes confirmed an outflow resistance distal to the trabecular meshwork (TM) that can be reduced. In an anterior segment model, nitric oxide increased the outflow facility even after the TM was removed [2]. Spectral-domain optical coherence tomography (SD-OCT) could detect outflow tract vessel dilation at the level of the collector channels (CC) and scleral vascular plexus (SVP) [2]. The results of these recent studies match historic ones that examined the impact of nitroglycerin [3–5] and hydralazine [3] on intraocular pressure (IOP) and outflow. Distal outflow resistance is pronounced in glaucoma patients. TM bypass [6, 7] and ablation [8–11] procedures were expected to reduce IOP to the level of episcleral venous pressure, around 8 mmHg [12].

Surprisingly, only approximately 0.3% of surgeries achieve this goal [11]. Moreover, a correlation of pre- and early postoperative IOP (before glaucoma medications are resumed) suggests that the baseline IOP reflects the severity of post-TM outflow pathology [8]. A glaucomatous change of the distal outflow tract that can cause such a significant resistance in small vessels can likely only be detected by high-resolution imaging techniques.

In our porcine eye model, we recently refined a benzyl alcohol benzyl benzoate (BABB) protocol as a clearing technique to overcome the opaqueness of the sclera and apply high-resolution imaging to the deep perilimbal structures. This requires staining the outflow vessels with lectin-labeled fluorophores and then using ribbon scanning confocal microscopy (RSCM) to reconstruct the outflow tract virtually [13]. The process of assembling several million confocal images is computation-intense but could be accomplished using a high-performance computer cluster [13, 14].

The human outflow tract has not been analyzed at such high resolution. Here, we apply this approach to two left eyes procured from octogenarian donors. The number of eyes had to be limited in this explorative study due to the intense computational requirements to process several terabytes of data in millions of confocal images. Canalogram studies of human eyes suggested a perilimbal, proximal vessel network with a density [15] similar to that of porcine eyes [16–19]. We hypothesized that an RSCM analysis of the lectin-labeled outflow tract of human eyes would match our findings in porcine eyes. Those studies found far more CCs than had been described before, with greater nasal CC openings, and a larger SVP in the inferior limbus

## Materials and methods

### Whole eye lectin perfusion

No human subjects were involved in the investigation. Human donor eye specimens were received from the Center for Organ Recovery and Education (CORE, Pittsburgh, PA). Only whole eyes without a history of glaucoma and procured within 48 hours of death were accepted. We analyzed a left eye from an 85-year-old female (eye 1) and a left eye from an 88-year-old male (eye 2). Eye 1 had no ocular history and eye 2 had a history of phacoemulsification with a posterior chamber intraocular lens implant. After perfusion with phosphate-buffered saline (PBS) for 30 minutes, a 300 µl intracameral bolus of a lectin-fluorophore conjugate (200 µg/mL) was applied, followed by constant-pressure infusion of the label at 20 µg/mL and 15 mmHg for 90 minutes. Eyes were stained with rhodamine-labeled lectin (#RLK-2200, Vector Laboratories, Inc, Burlingame, CA, USA). The eyes were subsequently fixed by perfusion with 4% paraformaldehyde (PFA) for 90 minutes and left in 4% PFA overnight, bisected along the equator, anterior segments cut into quadrants, and marked with indicator cuts according to their anatomical position. The resulting anterior segments were then cleared with BABB according to prior protocols [13].

### Image capture

Samples were imaged and processed as described before [13, 14]. Briefly, full-thickness perilimbal scans were acquired with a confocal microscope designed for high-speed ribbon-scanning and large scale image stitching (RS-G4, Caliber I.D., Andover, MA, USA). The system was fitted with a scanning stage (SCANplus IM 120 × 80, #00-24-579-0000; Märzhäuser Wetzlar GmbH & Co. KG, Wetzlar, Germany) and an Olympus 25×, 1.05 NA water immersion objective (XLPLN25XWMP2; Olympus). Volumetric scans were acquired with a voxel size of 0.365 × 0.365 × 2.43 μm. A scan-zoom of 1.5 was used during acquisition to achieve the desired resolution. Images were acquired over a single channel with an excitation wavelength of 561 nm and an emission filter of 630/60. Laser percentage, high voltage (HV), and offset were held constant throughout the volume at 7, 85, and 15, respectively.

### Image processing

The resulting series of Tagged Image File Format (TIFF) volumes comprised of 422 to 520 images were converted to the Bitplane Imaris IMS format. Each of the volumes contained between 548 and 889 gigabytes of data. To facilitate analysis, each of the IMS files was placed on a specialized high-speed solid-state file server equipped with 20 gigabits of network bandwidth. The IMS files were made accessible to multiple workstations equipped with a recent high-end Intel i7 processor and an NVIDIA GTX 1080 graphics card. All workstations were connected by 10-gigabit networking. We reconstructed the limbal outflow structures with image analysis software (Imaris 9.2, BitPlane AG, Zurich, Switzerland) and explored it in both 2D and 3D views. We identified the TM, Schlemm’s canal (SC), CCs, and SVPs with the surface function in Imaris. The range of grain size values used to resolve these structures was between 2.6 to 3.0 µm. Other parameters were a background subtraction with a diameter of the largest sphere fitting into the object of 10 to 200 µm, a background subtraction threshold value of 480 to 1387, and a filter to remove particles under 1.0×104 to 4.2×108 voxels. We created representative surfaces of the outflow tract structures up to 1000 µm from the distal TM. Surface creation parameters were refined to optimally resolve structural details of each region. Detailed Imaris surface creation parameters are provided in Supplementary Table 1. These surfaces were manually labeled as either TM/SC, CC, or SVP.

The TM were defined by a densely stained region of porous tissue at the base of each quadrant that appeared together with distinct beams. The beginnings of the CCs were defined by the appearance of a dark lumen surrounded by a bright wall with a beginning at the level of SC and a distal connection to the relatively superficial, perpendicularly oriented SVP. We observed different types of CC patterns of reaches at their catchment scale: meandering, braided, anabranching [20], and anastomosing. Given these differences, we decided to introduce the term “CC unit” as a region with a singular catchment area.

Using Imaris slice mode, we counted and measured the CC openings along the XY-plane at their sites of connection to the SC and SVP. The widest and narrowest lengths of these openings were recorded. All measurements were taken at the outermost point of proximal and distal confluence. Outlier CCs that continued beyond the 1,000 µm mark were reconstructed for visual representation.

Outlines of the projections of long ciliary nerves, known as Axenfeld loops [21–23], were identified in Imaris as dark structures not labeled with lectins. They were followed through slice mode to determine their path. We inverted the brightness values to highlight structures not labeled with lectins (Fig. 7D).

### Statistical analysis

The parameters of volume and location were obtained for TM and SC, CC, and SVP with the Imaris statistics function. CC opening lengths and widths were measured in Imaris slice mode. Location, CSA, and ellipticity for all CC openings were recorded. Data were analyzed by location for each eye. Data from human eyes in this study were compared to porcine eyes from our prior study [13]. Statistical tests were performed in Python 3.6.

## Results

Whole human eyes could be labeled and fully cleared in approximately seven days using our modified BABB protocol [13] (**Fig 1**). Unlike other species, human sclera mostly lacks pigmentation and became completely transparent when the refractive index of the normally opaque tissue matched that of BABB. This occurred during the final BABB step and took about 20 minutes. The eyes could be left for as long as 2 months in BABB without a notable difference in transparency or fluorescent signal. The scan time was 64 hours for eye 1 and 51 for eye 2, respectively. Lectin-labeled cleared anterior segments could be imaged via RSCM (**Fig. 2**). Lectin-labeled fluorophores intensely stained the entire outflow tract from the TM to the SVP. In eye 1, the SC was easy to delineate from the TM. However, SC in eye 2 could not be delineated from the TM as in eye 1. The conversion time from TIFF images to Imaris files was 6 hours for eye 1 and 8 hours for eye 2. The lectin binding pattern matched the one observed in our prior studies [13].

**Fig. 1:**
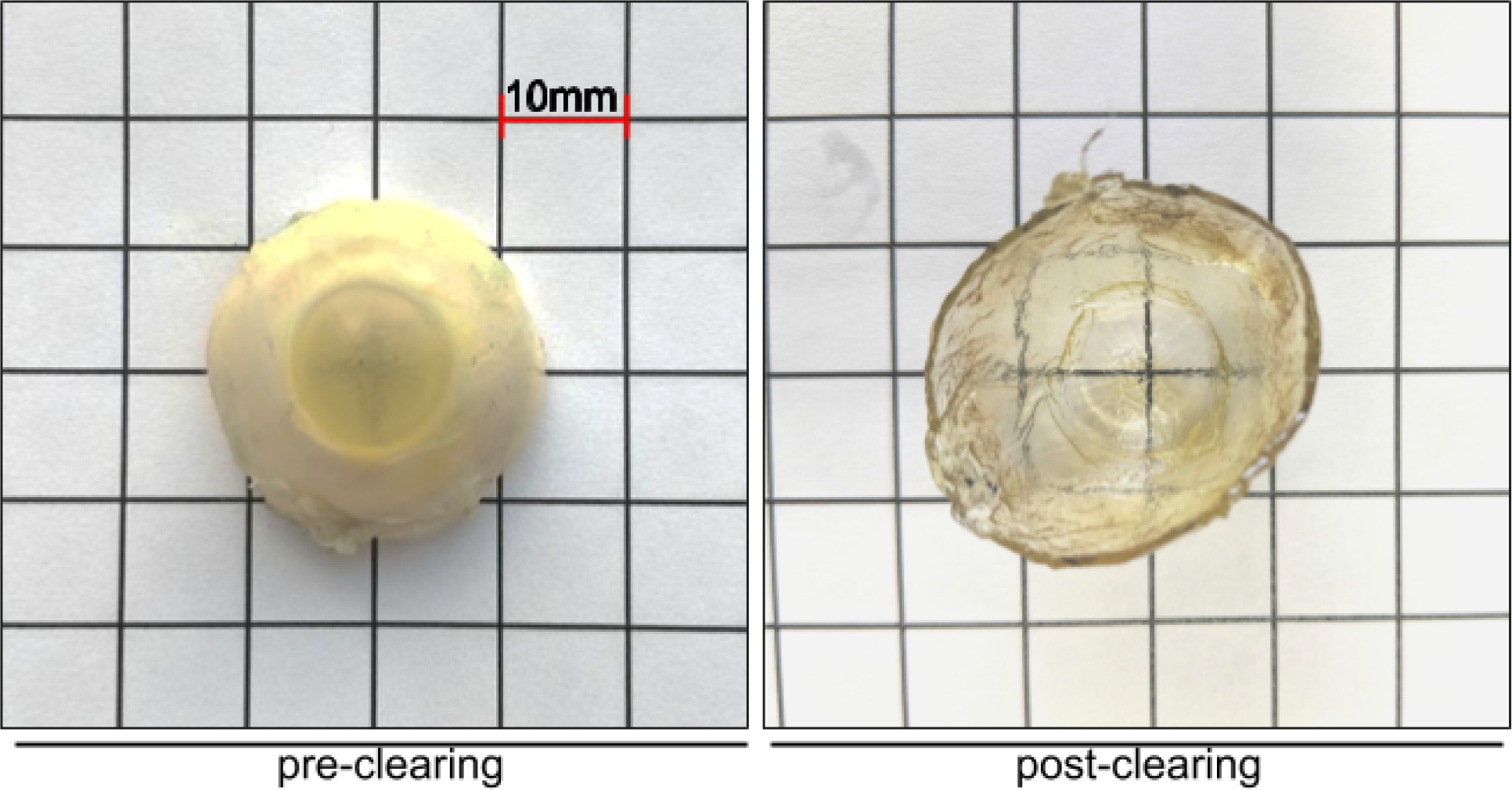
Macroscopic view of a BABB-cleared human eye. Human anterior segment pre and post BABB clearing.

**Figure 2:**
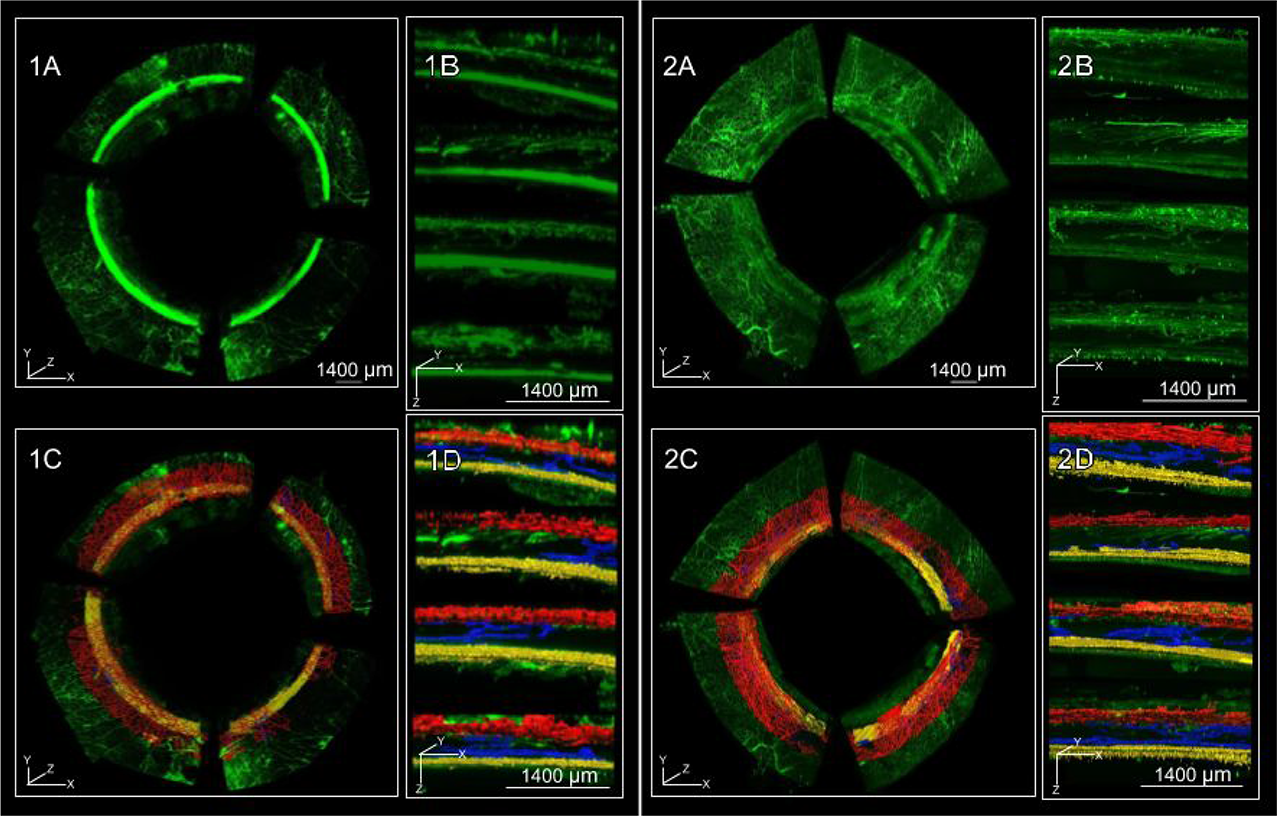
Volumetric limbal reconstruction by ribbon scanning confocal microscopy. RSCM reconstruction of eye 1 (1) and eye 2 (2) lectin-fluorophore perfused human eyes. Frontal (A) and sagittal (B) view of volumetric RSCM reconstructions. Frontal and sagittal views of labeled outflow tracts surfaces (C and D, respectively) with TM/SC in yellow, CC in blue, and SVP in red.

Computed volumes were displayed as radial charts (**Fig. 3**) according to the anatomic location in frontal view. Volume and region-dependent differences are presented in **Fig. 4**. The supranasal quadrant (SN_q_) of eye 1 had the largest SVP volume. The largest volume of TM and SC in this eye was in the infranasal quadrant (IN_q_). In eye 2, the largest SVP volume was in the infratemporal quadrant (IT_q_). The largest TM and SC volume in this eye was in the supratemporal quadrant (ST_q_). Some CC units had a wider area of catchment compared to others. Most of the CC units initiated from SC via several individual collectors before converging into a single lumen. In eye 1, 13 proximal CC openings connected to 20 distal CC openings through eight CC units. In eye 2, 13 proximal CC openings connected to 15 distal CC openings through 18 CC units. CC volume was greatest in the ST_q_. The lowest CC volume was found in the IN0_q_ of eye 1 and the IT_q_ of eye 2. In eye 1, CC volumes were about 13.3 times smaller than the SVP volumes, while in eye 2, CC volumes were 7.7 times smaller than SVP volumes. There were strikingly different CC morphologies that varied from quadrant to quadrant (**Fig. 5**).

**Figure 3:**
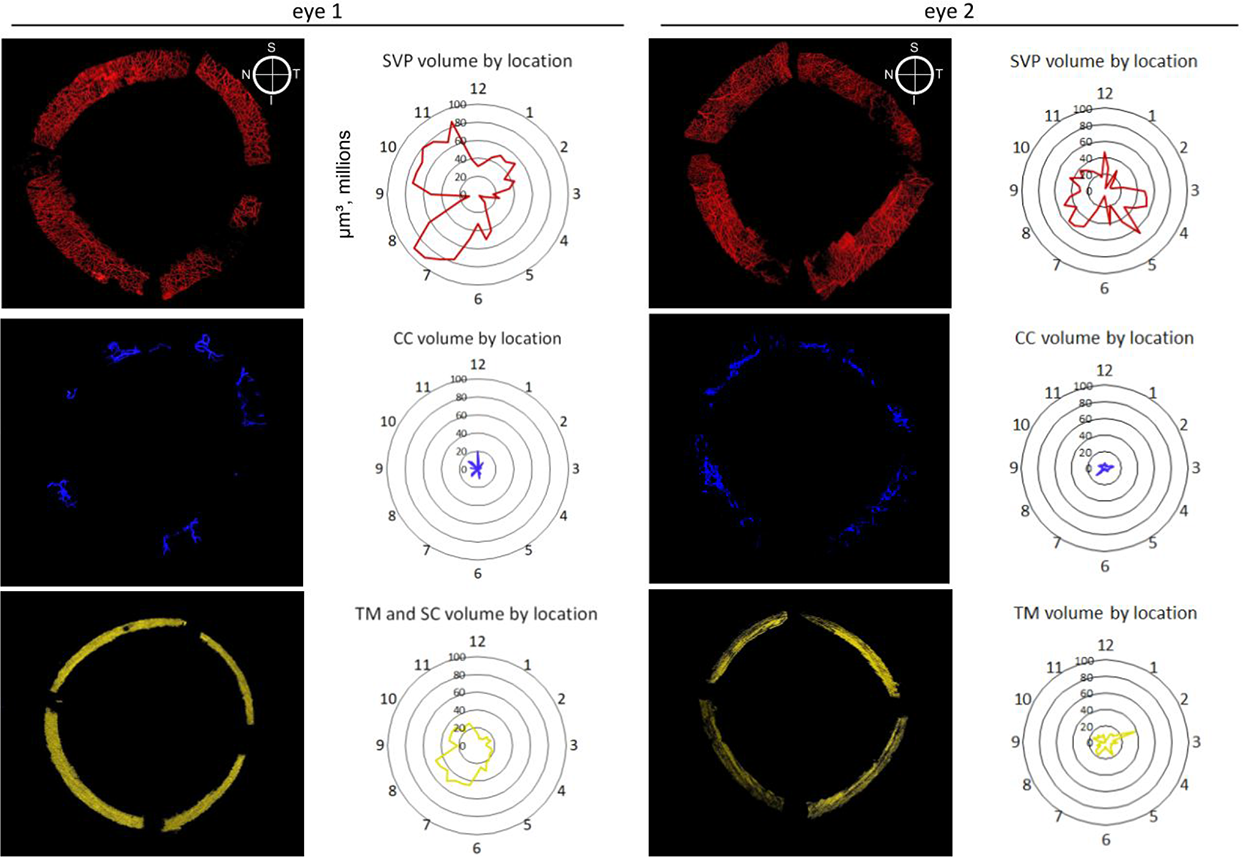
Radial charts with volumes. Frontal view of outflow tract structures. Volume measurements are plotted to correspond to their anatomical locations. TM/SC in yellow, CC in blue, SVP in red.

**Figure 4:**
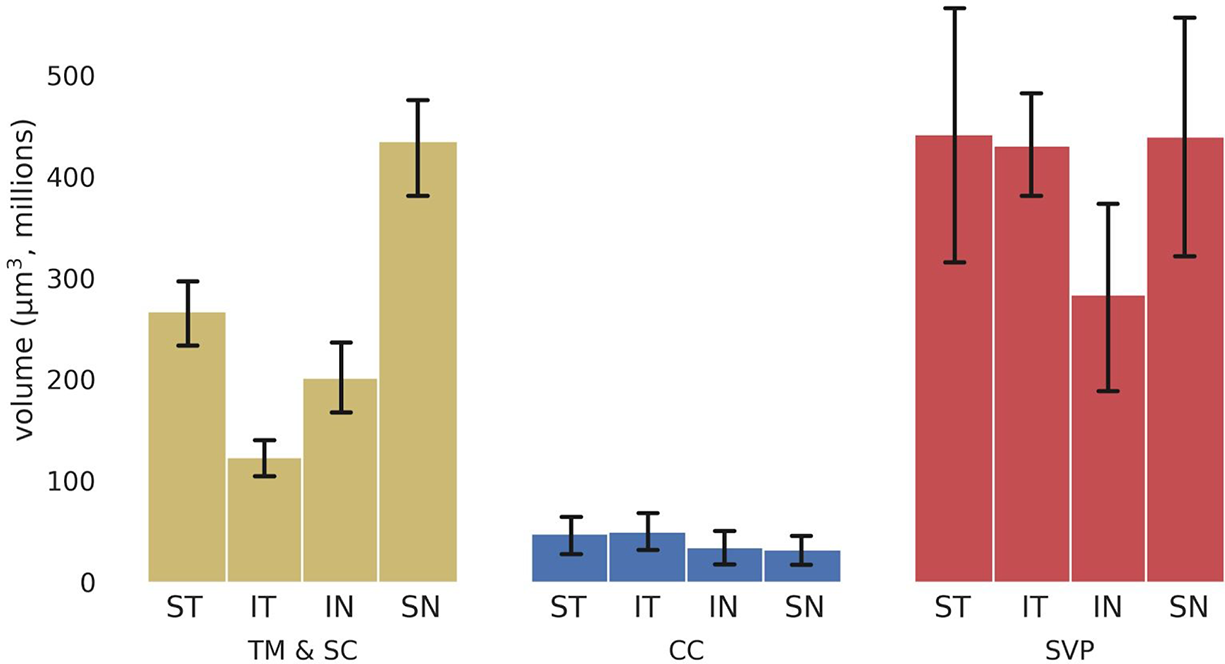
Outflow tract volumes. Volumes of outflow structures by anatomic location for eye 1 (top) and eye 2 (bottom) eye. SN: supranasal, ST: supratemporal, IT: infratemporal, IN: infranasal (averages with error bars using standard deviation).

**Fig. 5:**
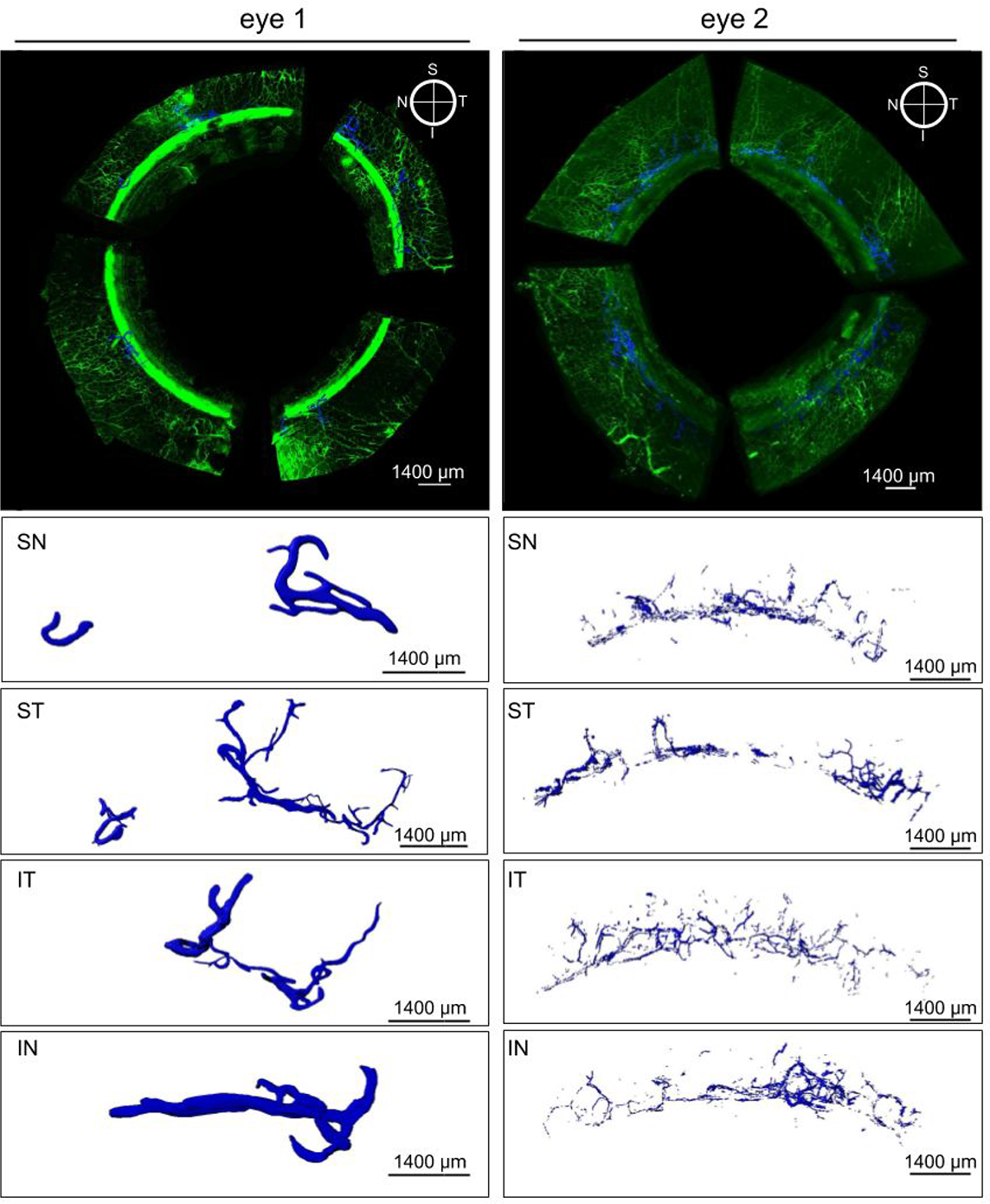
Collector channel morphology. Top panels show a volumetric reconstruction of eyes with CCs in blue. Bottom panels SN-IN show magnified CC morphology in each quadrant. SN: supranasal, ST: supratemporal, IT: infratemporal, IN: infranasal.

There were segmental differences in the CSAs of CC orifices at their proximal openings at the level of SC and at their distal points of confluence with the SVP (**Fig. 6A**). CC opening CSAs ranged from 9911.1µm^2^ to only 161.5µm^2^. Proximal openings were significantly larger than distal ones (P=0.017, 71.6%). The largest difference in average proximal and distal CSA were in the IT_q_ and the least difference in the ST_q_. The human CC opening CSA was significantly larger than the porcine CC opening CSA (p=7.37×10^-9^). The ellipticity of CC openings was compared at their proximal and distal ends. CC openings were more circular distally than proximally (1.42 and 1.18 times, respectively, P=0.0009) (**Fig 6B**). There was a significant linear relationship between CSA and cross-section circularity in both the human (P=1.88×10^-6^, r=-0.46) and porcine eye examined before [13] (P=0.0018, r=-0.21, linear regression) indicating that a large CSA is correlated with higher ellipticity or flatness.

**Fig. 6:**
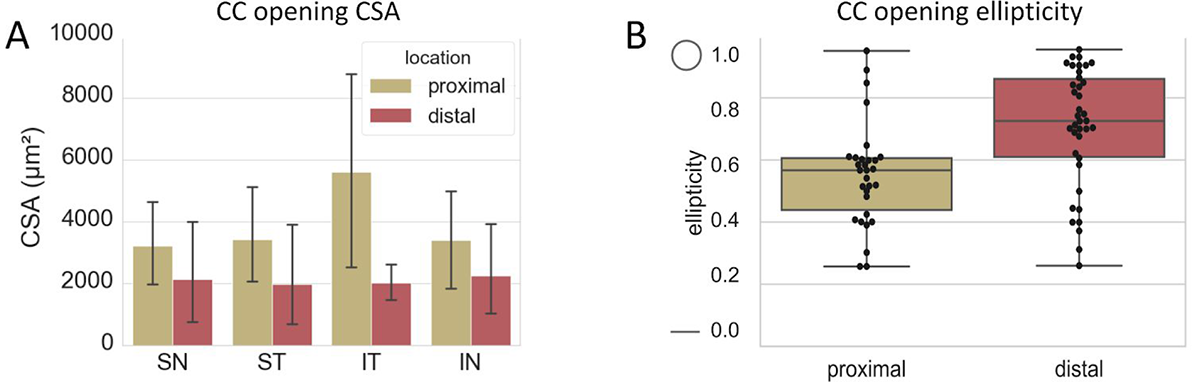
Cross-section areas and ellipticities of CC orifices at their proximal points of origin and at their distal points of connection with the SVP. A) Cross-section areas (CSA) of CC openings at their proximal points of connection with SC and their distal points of connection with the SVP. SN: supranasal, ST: supratemporal, IT: infratemporal, IN: infranasal. B) CC opening ellipticities of the proximal and distal ends. 1.0: perfect circle, 0.0: perfect line (averages with error bars using standard deviation).

**Fig. 7:**
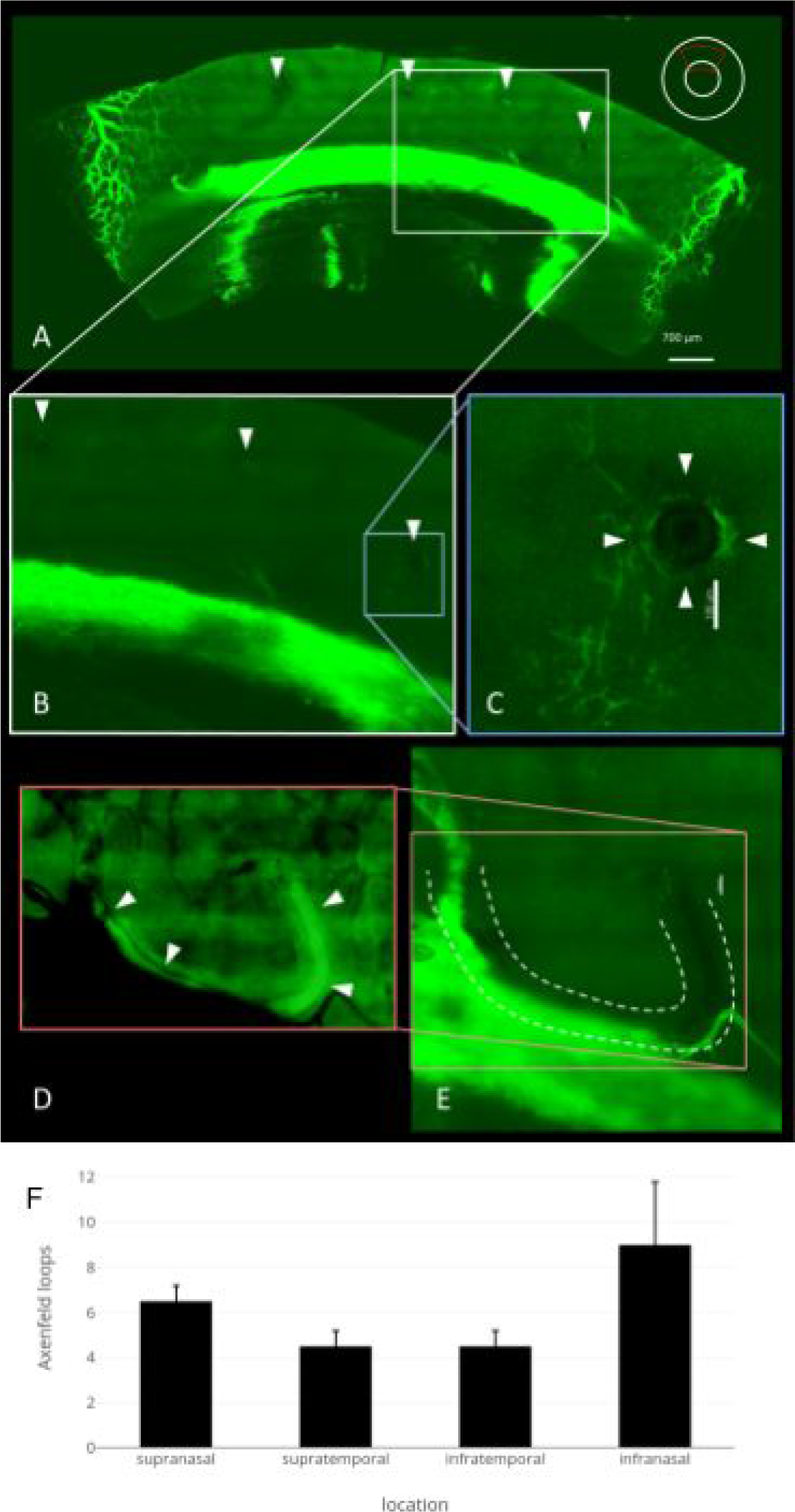
Axenfeld nerve loops could readily be identified in all quadrants. A) Upper quadrant of eye 2. Axenfeld loops are visible as dark structures void of lectin staining (white arrowheads). B) Magnified view of the area in A shows Axenfeld loop that enters the sclera. C) Magnified view of Axenfeld loop (white arrowheads) with perineural vasculature. D) The intrascleral path of the long ciliary nerve with Axenfeld loops can be more easily appreciated by image inversion. E) Direct proximity to the outer wall of Schlemm’s canal. F) Axenfeld loops per quadrant (averages with error bars using standard deviation).

The average CSA of individual CCs along their complete lengths in the human eye was 5221.1± 3521.1 µm^2^, greater than those in the porcine eye, 3374.0±2472.8 µm^2^ (P=0.002, 1.54 times greater, **Supplemental** **Fig. 1**). However, there was a lower CC density (0.52% coverage) in the human eye compared to the porcine eye (1.00% coverage, 1.92 times greater, P=0.014). In a representative sample of CC lengths measured, human CCs traveled 2533.4±1413.6 µm, a significantly longer distance than traveled by those in the porcine eye, 994.0±194.5 µm (P=0.004, 2.6 times longer, **Supplemental Fig. 1**). The relationship and connections of the elements of the conventional outflow system were assembled into a flythrough movie of eye 1 with all quadrants **(Supplemental material 2)** and at higher magnification the supranasal quadrant **(Supplemental material 3)**.

In eye 1, 27 Axenfeld loops were observed with an average of 6.8±2.3 per quadrant. In eye 2, 22 Axenfeld loops were seen with an average of 5.5±1.0 per quadrant. Most could be found in the IN_q_ followed by the SN_q_ (**Fig. 7**).

## Discussion

A growing body of evidence points to an outflow resistance distal to the TM as an essential aspect of both healthy [3,24,25] and glaucomatous [8–11] physiology. In this study, we provided a comprehensive overview of the human conventional outflow tract’s 3D architecture at an unprecedented resolution.

We analyzed variations in the microanatomy and catchment of CC units. We assessed the similarities and differences we found between the human and porcine outflow tract. Fluorophore-labeled lectins were able to stain the human outflow tract, validating the modified BABB protocol we developed in porcine eyes for imaging the limbus full-thickness at submicron confocal microscopy resolution [13]. Lectins are carbohydrate-binding proteins with a high specificity [26, 27] and can be used to study vessels by binding to their glycocalyx [28–30]. This new tool might be especially useful as Sienkiewicz et al.’s data [31] suggests that the accumulation of glycosylation end products in the TM may play a role in glaucoma. Although the current RSCM data acquisitions created 4-fold more data compared to our prior study, the process from staining to clearing and scanning was reasonably fast, taking approximately one week per eye, whereas a conventional confocal microscope would have required many months for the same acquisition. The combination of specialized toolsets to facilitate analysis, including purpose-built solid-state file servers and the Bitplane Imaris software suite, were critical to improving handling. These methods provide a new tool to researchers working on deciphering the aspects of structure and function of the outflow tract at high 3D resolution.

We observed significant inter-quadrant and inter-individual variations in the microanatomy of the outflow tract through volumetric reconstruction of RSCM-derived images. Our findings match Bentley et al. who found that proximal CC openings varied from simple and ellipsoid to tethered flaps and bridges [32]. In the future, fluid dynamics studies that use RSCM data-based mesh surfaces will examine the specific roles and impacts of these different CC unit morphologies.

The importance and potential vulnerability of CCs are highlighted by the fact that their total volume is only about 10% of the volume of the SVP. Depending on the compliance of CCs and the surrounding tissue, it is possible that the highly elliptical shape of larger CSA vessels may imply that the tract’s capacity for fluid transport has not been reached. With higher local pressures, elliptical cross-sections may be forced into a fully turgid circular shape, as already seen in the smaller distal CC openings, to act like pressure valves that maintain IOP at a set minimum value.

In comparison to the RSCM analysis of BABB-cleared porcine eyes [13], we found that CCs in human eyes were only about half of the number of porcine eyes and traveled over twice the distance from Schlemm’s canal to the SVP. These results indicate that human eyes not only have far fewer collector channels but that their anatomy differs fundamentally by spanning several clock hours and having a 50% greater CSA. This came as a surprise because recent lower resolution canalogram studies of perilimbal outflow structures suggested a density and function of outflow vessels in human eyes [15] similar to that in porcine eyes [16–19]. The smaller number, longer course, and flatter, collapsible nature of CCs in human eyes may increase the risk of outflow failure and IOP elevation in glaucoma. Consistent with this, Hann et al. described 3.7 times more CC occlusions in eyes with glaucoma [33]. It will be interesting to compare the outflow vessel glycocalyx pattern in RSCM scans of eyes with glaucoma to look for clues of an altered wall adherence.

In the ST_q_ of eye 2, there was one region with a weak signal and low volumes that corresponded to the corneoscleral tunnel created during cataract surgery while the adjacent ones were enlarged. Considering that eye 2 had surgery and no history of glaucoma, it is tempting to speculate that the opening CSAs or the remaining adjacent vessels enlarged to facilitate flow. This matches an increased count of CCs at higher IOPs as reported by Hann et al. [33].

In the anterior segment, the long ciliary nerve has prominent branches of unknown function, termed Axenfeld loops [22]. Reese described them as a retroverted nerve loop positioned on the scleral surface, with occasionally accompanying cyst-like structures that are more prominent in glaucoma [23]. The fact that we observed 6 such loops per quadrant at regular intervals in proximity and parallel to CCs suggests that they might play a role related to the innervation of TM, scleral spur [34], collector channels or EVP [35] to participate in the autonomous regulation of outflow [36].

A limitation of our study is that a more comprehensive analysis with additional eyes would have been desirable, but the relative scarcity of human donor eyes and the intense computational demand required a focused approach.

In summary, we analyzed the human conventional outflow tract of octogenarian eyes in full-thickness, high-resolution and 3D. Using their CC glycosaminoglycan lectin staining profile, we characterized CSA, length, and branching patterns. Key differences between the human and porcine outflow tract may help explain why humans — but not pigs — frequently develop glaucoma. Future studies will need to assess glaucoma specific changes of morphology and glycocalyx composition of the conventional outflow tract.

## Acknowledgments

This study was funded by the Initiative to Cure Glaucoma of the Eye and Ear Foundation of Pittsburgh (NAL), by NEI Grant K08EY022737, by NIH CORE Grant P30 EY08098 to the Department of Ophthalmology, and an unrestricted grant from Research to Prevent Blindness, New York, NY.

## Supplemental material 2

Fly-through movie

Supplemental material 2: Fly-through movie of eye 1.

**Supplemental figure 1:**
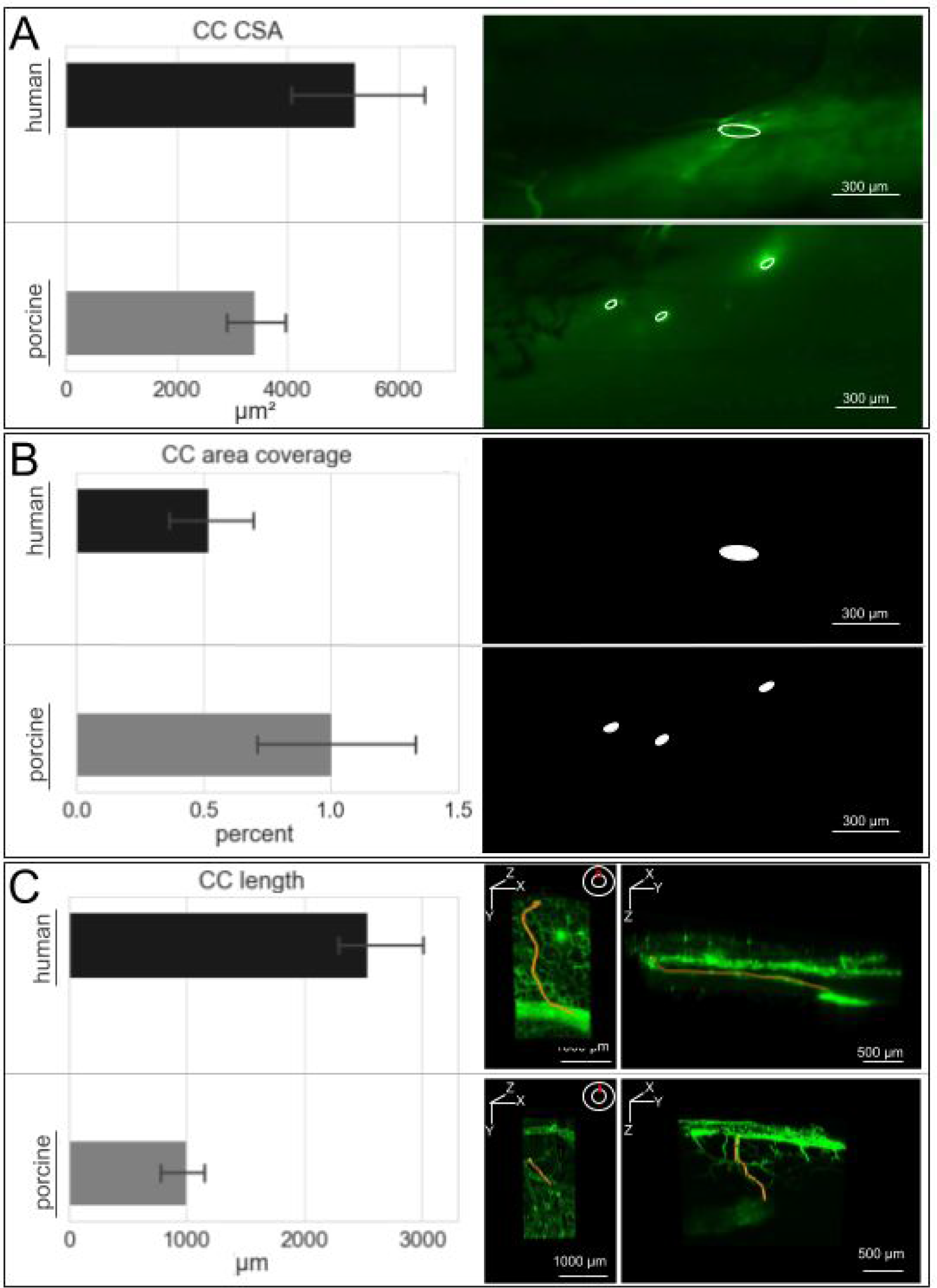
A) Human and porcine CC CSA (left) and representative images of measured CC openings (top: human, bottom: porcine). B) Human and porcine CC area coverage (left) and images of CC opening areas as shown in A. C) Human and porcine CC length (left) and representative CCs traced (orange). CC: collector channel, CSA: cross-section area

## Supplemental material 3

Quadrant B sped up to 2X

Supplemental material 3: Fly-through movie of the supranasal quadrant of eye 1.

## Supplemental table 1

### Surface Parameter Ranges

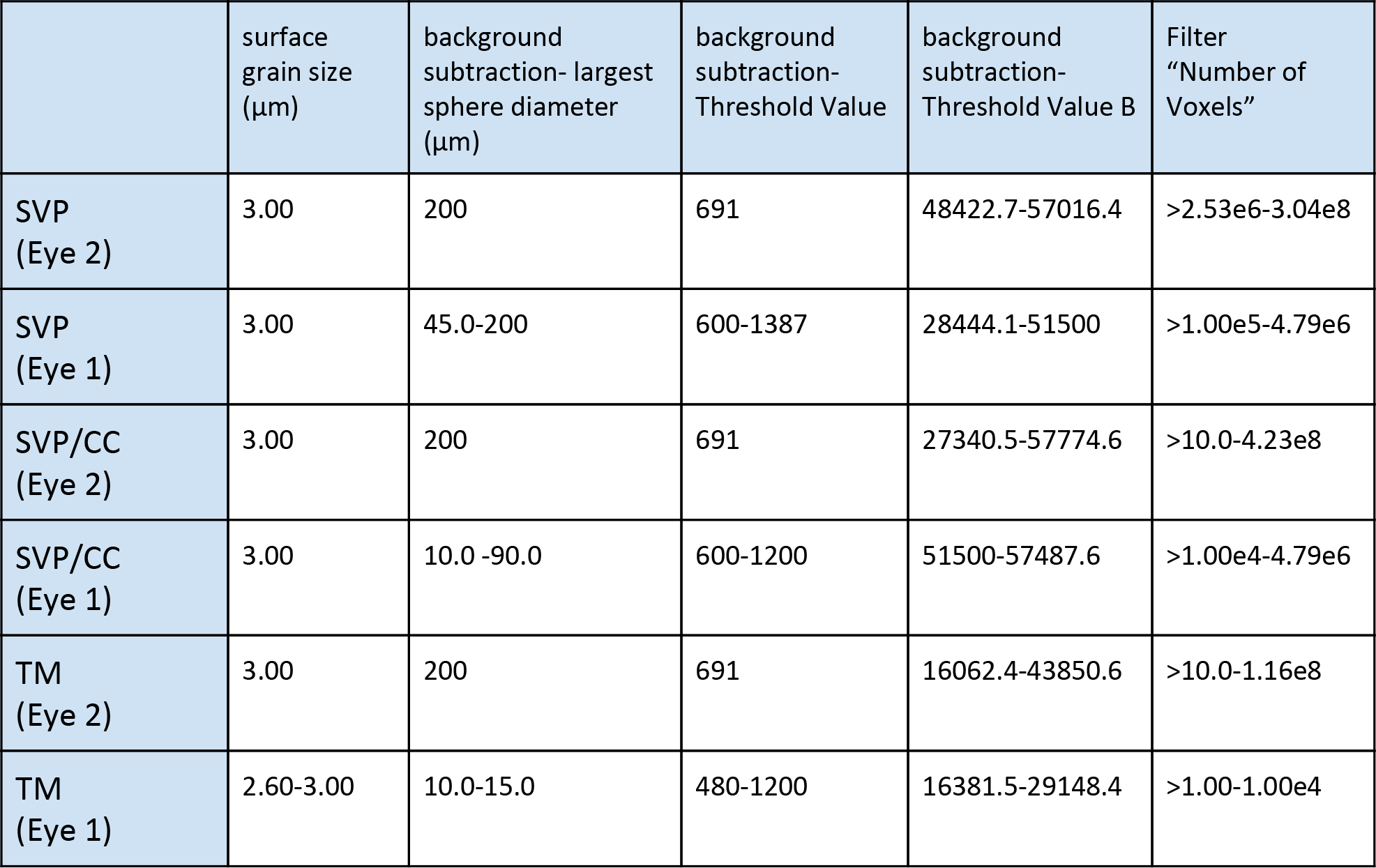

|  | surface grain size (μm) | background subtraction- largest sphere diameter (μm) | background subtraction- Threshold Value | background subtraction- Threshold Value B | Filter "Number of Voxels" |
| --- | --- | --- | --- | --- | --- |
| SVP (Eye 2) | 3.00 | 200 | 691 | 48422.7-57016.4 | >2.53e6-3.04e8 |
| SVP (Eye 1) | 3.00 | 45.0-200 | 600-1387 | 28444.1-51500 | >1.00e5-4.79e6 |
| SVP/CC (Eye 2) | 3.00 | 200 | 691 | 27340.5-57774.6 | >10.0-4.23e8 |
| SVP/CC (Eye 1) | 3.00 | 10.0 -90.0 | 600-1200 | 51500-57487.6 | >1.00e4-4.79e6 |
| TM (Eye 2) | 3.00 | 200 | 691 | 16062.4-43850.6 | >10.0-1.16e8 |
| TM (Eye 1) | 2.60-3.00 | 10.0-15.0 | 480-1200 | 16381.5-29148.4 | >1.00-1.00e4 |

## Eye 2

| **SVP** | surface grain size | background subtraction- largest sphere diameter | background subtraction- Threshold Value | background subtraction- Threshold Value B | Filter "Number of Voxels" |
| --- | --- | --- | --- | --- | --- |
| Q4, Surface 13 | 3.00 µm | 200 µm | 691 | 48422.7 | > 3.48e7 |
| Q3, Surface 5 | 3.00 µm | 200 µm | 691 | 57016.4 | > 2.53e6 |
| Q2, Surface 29 | 3.00 µm | 200 µm | 691 | 51045.5 | > 4.44e6 |
| Q1, Surface 26 | 3.00 µm | 200 µm | 691 | 52597.5 | > 3.04e8 |
| Eye 2 SVP Range | 3.00 µm | 200 µm | 691 | 48422.7-57016.4 | >2.53e6-3.04e8 |

| **SVP/CC** | surface grain size | background subtraction- largest sphere diameter | background subtraction- Threshold Value | background subtraction- Threshold Value B | Filter "Number of Voxels" |
| --- | --- | --- | --- | --- | --- |
| Q4, Surface 15 | 3.00 µm | 200 µm | 691 | 57774.6 | >4.77e6 |
| Q4, Surface 14 | 3.00 µm | 200 µm | 691 | 35586.6 | >4.73e7 |
| Q3, Surface 13 | 3.00 µm | 200 µm | 691 | 27340.5 | >10.0 |
| Q1, Surface 25 | 3.00 µm | 200 µm | 691 | 50990.8 | >2.59e6<br><4.23e8 |
| Eye 2 SVP/CC Range | 3.00 µm | 200 µm | 691 | 27340.5-57774.6 | >10.0-4.23e8 |

| **TM** | surface grain size | background subtraction- largest sphere diameter | background subtraction- Threshold Value | background subtraction- Threshold Value B | Filter "Number of Voxels" |
| --- | --- | --- | --- | --- | --- |
| Q4, Surface 2 | 3.00 µm | 200 µm | 691 | 36092.5 | >5.22e6 |
| Q4, Surface 11 | 3.00 µm | 200 µm | 691 | 16062.4 | >7.74e7 |
| Q2, Surface 7 | 3.00 µm | 200 µm | 691 | 43850.6 | >2.20e7<br><1.16e8 |
| Q1, Surface 14 | 3.00 µm | 200 µm | 691 | 22287 | >10.0 |
| Eye 2 TM Range | 3.00 µm | 200 µm | 691 | 16062.4-43850.6 | >10.0-1.16e8 |

## Eye 1

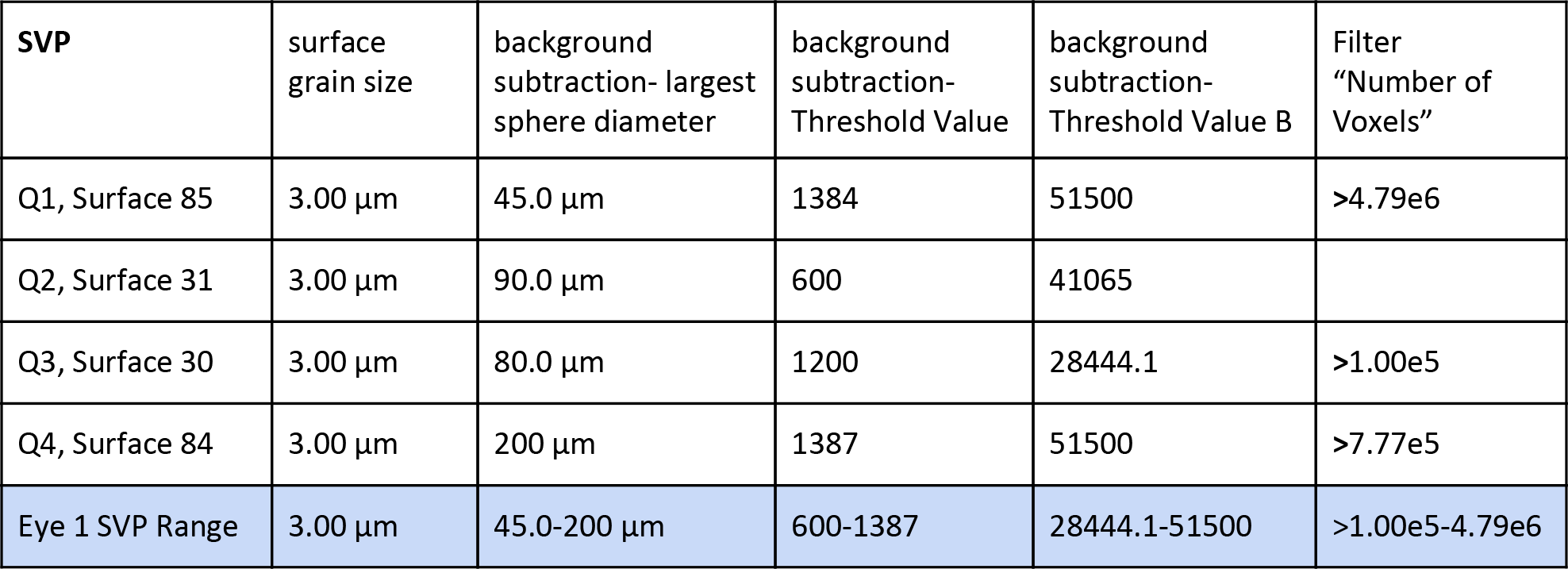
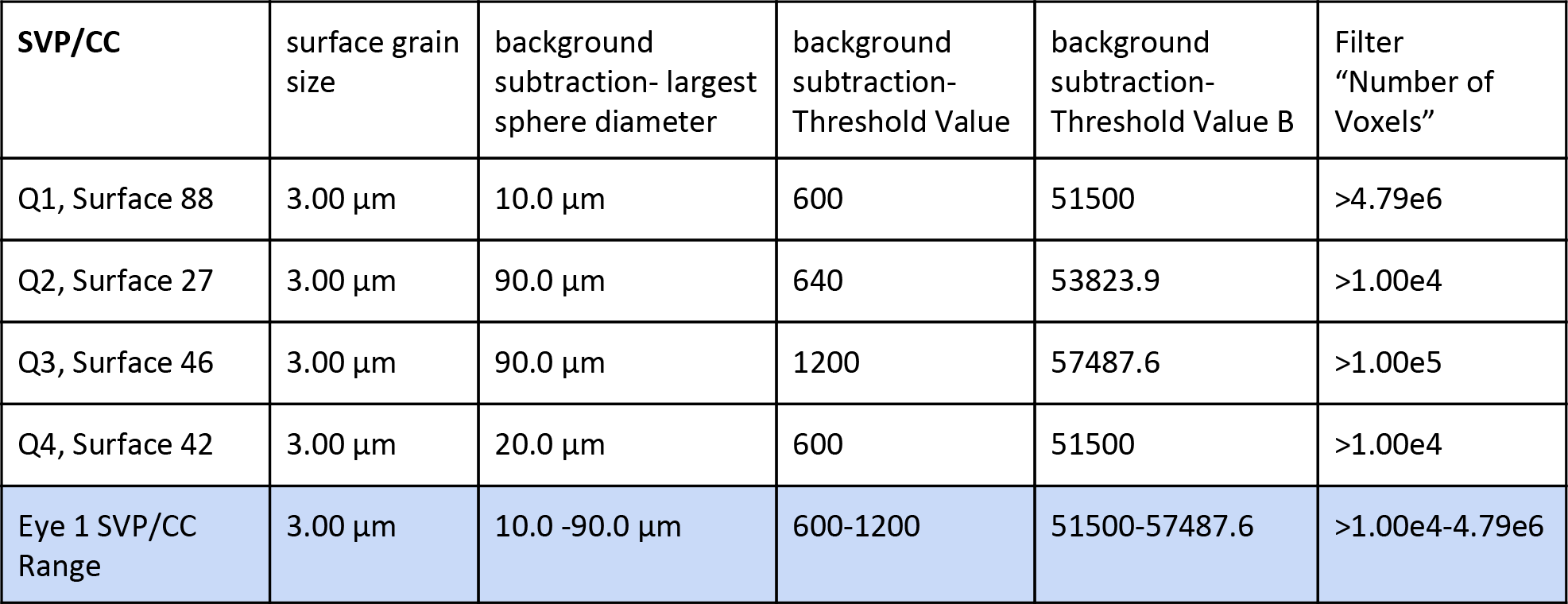
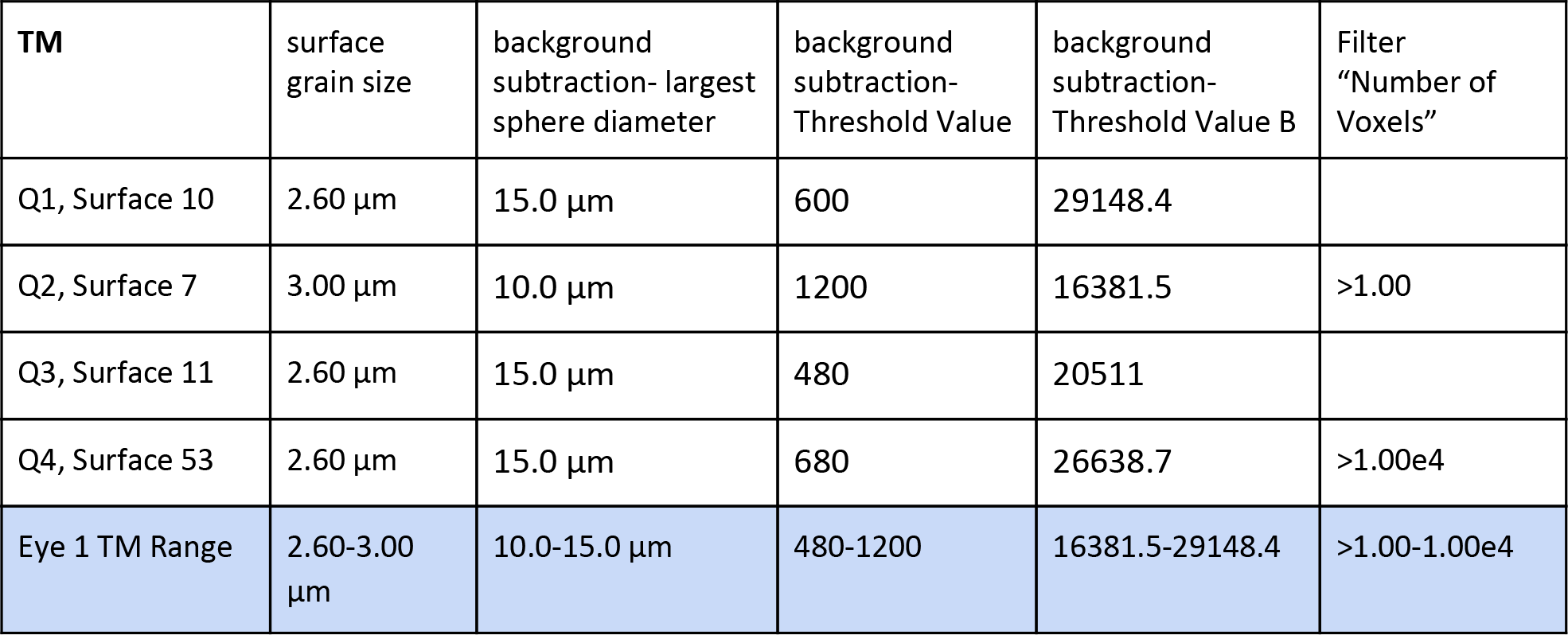

| **SVP** | surface grain size | background subtraction- largest sphere diameter | background subtraction- Threshold Value | background subtraction- Threshold Value B | Filter "Number of Voxels" |
| --- | --- | --- | --- | --- | --- |
| Q1, Surface 85 | 3.00 µm | 45.0 µm | 1384 | 51500 | >4.79e6 |
| Q2, Surface 31 | 3.00 µm | 90.0 µm | 600 | 41065 |  |
| Q3, Surface 30 | 3.00 µm | 80.0 µm | 1200 | 28444.1 | >1.00e5 |
| Q4, Surface 84 | 3.00 µm | 200 µm | 1387 | 51500 | >7.77e5 |
| Eye 1 SVP Range | 3.00 µm | 45.0-200 µm | 600-1387 | 28444.1-51500 | >1.00e5-4.79e6 |

| **SVP/CC** | surface grain size | background subtraction- largest sphere diameter | background subtraction- Threshold Value | background subtraction- Threshold Value B | Filter "Number of Voxels" |
| --- | --- | --- | --- | --- | --- |
| Q1, Surface 88 | 3.00 µm | 10.0 µm | 600 | 51500 | >4.79e6 |
| Q2, Surface 27 | 3.00 µm | 90.0 µm | 640 | 53823.9 | >1.00e4 |
| Q3, Surface 46 | 3.00 µm | 90.0 µm | 1200 | 57487.6 | >1.00e5 |
| Q4, Surface 42 | 3.00 µm | 20.0 µm | 600 | 51500 | >1.00e4 |
| Eye 1 SVP/CC Range | 3.00 µm | 10.0 -90.0 µm | 600-1200 | 51500-57487.6 | >1.00e4-4.79e6 |

| **TM** | surface grain size | background subtraction- largest sphere diameter | background subtraction- Threshold Value | background subtraction- Threshold Value B | Filter "Number of Voxels" |
| --- | --- | --- | --- | --- | --- |
| Q1, Surface 10 | 2.60 µm | 15.0 µm | 600 | 29148.4 |  |
| Q2, Surface 7 | 3.00 µm | 10.0 µm | 1200 | 16381.5 | >1.00 |
| Q3, Surface 11 | 2.60 µm | 15.0 µm | 480 | 20511 |  |
| Q4, Surface 53 | 2.60 µm | 15.0 µm | 680 | 26638.7 | >1.00e4 |
| Eye 1 TM Range | 2.60-3.00 µm | 10.0-15.0 µm | 480-1200 | 16381.5-29148.4 | >1.00-1.00e4 |

## Eye 2

### SVP

- Eye 2, Quadrant 4, Surface 13
Surface Grain Size = 3.00 µm
Enable Eliminate Background = true
Diameter Of Largest Sphere = 200 µm
[Threshold]
Enable Automatic Threshold = false
Manual Threshold Value = 691
Active Threshold = true
Enable Automatic Threshold B = true
Manual Threshold Value B = 48422.7
Active Threshold B = false
[Classify Surfaces]
“Number of Voxels” above 3.48e7

- Eye 2, Quadrant 3, Surface 5
Surface Grain Size = 3.00 µm
Enable Eliminate Background = true
Diameter Of Largest Sphere = 200 µm
[Threshold]
Enable Automatic Threshold = false
Manual Threshold Value = 691
Active Threshold = true
Enable Automatic Threshold B = true
Manual Threshold Value B = 57016.4
Active Threshold B = false
[Classify Surfaces]
“Number of Voxels” above 2.53e6

- Eye 2, Quadrant 2, Surface 29
Surface Grain Size = 3.00 µm
Enable Eliminate Background = true
Diameter Of Largest Sphere = 200 µm
[Threshold]
Enable Automatic Threshold = false
Manual Threshold Value = 691
Active Threshold = true
Enable Automatic Threshold B = true
Manual Threshold Value B = 51045.5
Active Threshold B = false
[Classify Surfaces]
“Number of Voxels” above 4.44e6

- Eye 2, Quadrant 1, Surface 26
Surface Grain Size = 3.00 µm
Enable Eliminate Background = true
Diameter Of Largest Sphere = 200 µm
[Threshold]
Enable Automatic Threshold = false
Manual Threshold Value = 691
Active Threshold = true
Enable Automatic Threshold B = true
Manual Threshold Value B = 52597.5
Active Threshold B = false
[Classify Surfaces]
“Number of Voxels” above 3.04e8

## SVP/CC

- Eye 2, Quadrant 4, Surface 15
Surface Grain Size = 3.00 µm
Enable Eliminate Background = true
Diameter Of Largest Sphere = 200 µm
[Threshold]
Enable Automatic Threshold = false
Manual Threshold Value = 691
Active Threshold = true
Enable Automatic Threshold B = true
Manual Threshold Value B = 57774.6
Active Threshold B = false
[Classify Surfaces]
“Number of Voxels” above 4.77e6

- Eye 2, Quadrant 4, Surface 14
Surface Grain Size = 3.00 µm
Enable Eliminate Background = true
Diameter Of Largest Sphere = 200 µm
[Threshold]
Enable Automatic Threshold = false
Manual Threshold Value = 691
Active Threshold = true
Enable Automatic Threshold B = true
Manual Threshold Value B = 35586.6
Active Threshold B = false
[Classify Surfaces]
“Number of Voxels” above 4.73e7

- Eye 2, Quadrant 3, Surface 13
Surface Grain Size = 3.00 µm
Enable Eliminate Background = true
Diameter Of Largest Sphere = 200 µm
[Threshold]
Enable Automatic Threshold = false
Manual Threshold Value = 691
Active Threshold = true
Enable Automatic Threshold B = true
Manual Threshold Value B = 27340.5
Active Threshold B = false
[Classify Surfaces]
“Number of Voxels” above 10.0

- Eye 2, Quadrant 1, Surface 25
Surface Grain Size = 3.00 µm
Enable Eliminate Background = true
Diameter Of Largest Sphere = 200 µm
[Threshold]
Enable Automatic Threshold = false
Manual Threshold Value = 691
Active Threshold = true
Enable Automatic Threshold B = true
Manual Threshold Value B = 50990.8
Active Threshold B = false
[Classify Surfaces]
“Number of Voxels” between 2.59e6 and 4.23e8

## TM

- Eye 2, Quadrant 4, Surface 2
Surface Grain Size = 3.00 µm
Enable Eliminate Background = true
Diameter Of Largest Sphere = 200 µm
[Threshold]
Enable Automatic Threshold = false
Manual Threshold Value = 691
Active Threshold = true
Enable Automatic Threshold B = true
Manual Threshold Value B = 36092.5
Active Threshold B = false
[Classify Surfaces]
“Number of Voxels” above 5.22e6

- Eye 2, Quadrant 4, Surface 11
Surface Grain Size = 3.00 µm
Enable Eliminate Background = true
Diameter Of Largest Sphere = 200 µm
[Threshold]
Enable Automatic Threshold = false
Manual Threshold Value = 691
Active Threshold = true
Enable Automatic Threshold B = true
Manual Threshold Value B = 16062.4
Active Threshold B = false
[Classify Surfaces]
“Number of Voxels” above 7.74e7

- Eye 2, Quadrant 2, Surface 7
Surface Grain Size = 3.00 µm
Enable Eliminate Background = true
Diameter Of Largest Sphere = 200 µm
[Threshold]
Enable Automatic Threshold = false
Manual Threshold Value = 691
Active Threshold = true
Enable Automatic Threshold B = true
Manual Threshold Value B = 43850.6
Active Threshold B = false
[Classify Surfaces]
“Number of Voxels” between 2.20e7 and 1.16e8

- Eye 2, Quadrant 1, Surface 14
Surface Grain Size = 3.00 µm
Enable Eliminate Background = true
Diameter Of Largest Sphere = 200 µm
[Threshold]
Enable Automatic Threshold = false
Manual Threshold Value = 691
Active Threshold = true
Enable Automatic Threshold B = true
Manual Threshold Value B = 22287
Active Threshold B = false
[Classify Surfaces]
“Number of Voxels” above 10.0

## Eye 1

### SVP

- Eye 1, Quadrant 1, Surface 85
Surface Grain Size = 3.00 µm
Enable Eliminate Background = true
Diameter Of Largest Sphere = 45.0 µm
[Threshold]
Enable Automatic Threshold = false
Manual Threshold Value = 1384
Active Threshold = true
Enable Automatic Threshold B = false
Manual Threshold Value B = 51500
Active Threshold B = true
[Classify Surfaces]
“Number of Voxels Img=1” above 4.79e6

- Eye 1, Quadrant 2, Surface 31
Surface Grain Size = 3.00 µm
Enable Eliminate Background = true
Diameter Of Largest Sphere = 90.0 µm
[Threshold]
Enable Automatic Threshold = false
Manual Threshold Value = 600
Active Threshold = true
Enable Automatic Threshold B = true
Manual Threshold Value B = 41065
Active Threshold B = false
[Classify Surfaces]
“Volume” above 1.00e4 µm^3

- Eye 1, Quadrant 3, Surface 30
Surface Grain Size = 3.00 µm
Enable Eliminate Background = true
Diameter Of Largest Sphere = 80.0 µm
[Threshold]
Enable Automatic Threshold = false
Manual Threshold Value = 1200
Active Threshold = true
Enable Automatic Threshold B = false
Manual Threshold Value B = 28444.1
Active Threshold B = false
[Classify Surfaces]
“Number of Voxels Img=1” above 1.00e5

- Eye 1, Quadrant 4, Surface 84
Surface Grain Size = 3.00 µm
Enable Eliminate Background = true
Diameter Of Largest Sphere = 200 µm
[Threshold]
Enable Automatic Threshold = false
Manual Threshold Value = 1387
Active Threshold = true
Enable Automatic Threshold B = false
Manual Threshold Value B = 51500
Active Threshold B = true
[Classify Surfaces]
“Number of Voxels Img=1” above 7.77e5

## SVP/CC

- Eye 1, Quadrant 1, Surface 88
Surface Grain Size = 3.00 µm
Enable Eliminate Background = true
Diameter Of Largest Sphere = 10.0 µm
[Threshold]
Enable Automatic Threshold = false
Manual Threshold Value = 600
Active Threshold = true
Enable Automatic Threshold B = false
Manual Threshold Value B = 51500
Active Threshold B = true
[Classify Surfaces]
“Number of Voxels Img=1” above 4.79e6

- Eye 1, Quadrant 2, Surface 27
Surface Grain Size = 3.00 µm
Enable Eliminate Background = true
Diameter Of Largest Sphere = 90.0 µm
[Threshold]
Enable Automatic Threshold = false
Manual Threshold Value = 640
Active Threshold = true
Enable Automatic Threshold B = true
Manual Threshold Value B = 53823.9
Active Threshold B = false
[Classify Surfaces]
“Number of Voxels Img=1” above 1.00e4

- Eye 1, Quadrant 3, Surface 46
Surface Grain Size = 3.00 µm
Enable Eliminate Background = true
Diameter Of Largest Sphere = 90.0 µm
[Threshold]
Enable Automatic Threshold = false
Manual Threshold Value = 1200
Active Threshold = true
Enable Automatic Threshold B = false
Manual Threshold Value B = 57487.6
Active Threshold B = false
[Classify Surfaces]
“Number of Voxels Img=1” above 1.00e5

- Eye 1, Quadrant 4, Surface 42
Surface Grain Size = 3.00 µm
Enable Eliminate Background = true
Diameter Of Largest Sphere = 20.0 µm
[Threshold]
Enable Automatic Threshold = false
Manual Threshold Value = 600
Active Threshold = true
Enable Automatic Threshold B = false
Manual Threshold Value B = 51500
Active Threshold B = true
[Classify Surfaces]
“Number of Voxels Img=1” above 1.00e4

## TM

- Eye 1, Quadrant 1, Surface 10
Surface Grain Size = 2.60 µm
Enable Eliminate Background = true
Diameter Of Largest Sphere = 15.0 µm
[Threshold]
Enable Automatic Threshold = false
Manual Threshold Value = 600
Active Threshold = true
Enable Automatic Threshold B = false
Manual Threshold Value B = 29148.4
Active Threshold B = false
[Classify Surfaces]
“Volume” above 1.00e4 µm^3

- Eye 1, Quadrant 2, Surface 7

## References

1. McDonnell F, Dismuke WM, Overby DR, Stamer WD. Pharmacological regulation of outflow resistance distal to Schlemm’s canal. Am J Physiol Cell Physiol. 2018;315: C44–C51.

2. Waxman S, Wang C, Dang Y, Hong Y, Esfandiari H, Shah P, et al. Structure-Function Changes of the Porcine Distal Outflow Tract in Response to Nitric Oxide. Invest Ophthalmol Vis Sci. 2018;59: 4886–4895.

3. Schuman JS, Erickson K, Nathanson JA. Nitrovasodilator effects on intraocular pressure and outflow facility in monkeys. Exp Eye Res. 1994;58: 99–105.

4. Nathanson JA. Guanylyl cyclase activators: effects on ciliary process cyclic GMP metabolism and intraocular pressure. Invest Ophthalmol Vis Sci. 1988;29: 5.

5. Nathanson JA. Nitrovasodilators as a new class of ocular hypotensive agents. J Pharmacol Exp Ther. 1992;260: 956–965.

6. Fernández-Barrientos Y, García-Feijoó J, Martínez-de-la-Casa JM, Pablo LE, Fernández-Pérez C, García Sánchez J. Fluorophotometric study of the effect of the glaukos trabecular microbypass stent on aqueous humor dynamics. Invest Ophthalmol Vis Sci. 2010;51: 3327–3332.

7. Samuelson TW, Katz LJ, Wells JM, Duh Y-J, Giamporcaro JE, US iStent Study Group. Randomized evaluation of the trabecular micro-bypass stent with phacoemulsification in patients with glaucoma and cataract. Ophthalmology. 2011;118: 459–467.

8. Bussel II, Kaplowitz K, Schuman JS, Loewen NA, Trabectome Study Group. Outcomes of ab interno trabeculectomy with the trabectome by degree of angle opening. Br J Ophthalmol. 2015;99: 914–919.

9. Loewen RT, Roy P, Parikh HA, Dang Y, Schuman JS, Loewen NA. Impact of a Glaucoma Severity Index on Results of Trabectome Surgery: Larger Pressure Reduction in More Severe Glaucoma. PLoS One. 2016;11: e0151926.

10. Bussel II, Kaplowitz K, Schuman JS, Loewen NA, Group TS, Others. Outcomes of ab interno trabeculectomy with the trabectome after failed trabeculectomy. Br J Ophthalmol. 2014;99: 258–262.

11. Parikh HA, Bussel II, Schuman JS, Brown EN, Loewen NA. Coarsened Exact Matching of Phaco-Trabectome to Trabectome in Phakic Patients: Lack of Additional Pressure Reduction from Phacoemulsification. PLoS One. 2016;11: e0149384.

12. Sit AJ, McLaren JW. Measurement of episcleral venous pressure. Exp Eye Res. 2011;93: 291–298.

13. Waxman S, Loewen RT, Dang Y, Watkins SC, Watson AM, Loewen NA. High-Resolution, Three-Dimensional Reconstruction of the Outflow Tract Demonstrates Segmental Differences in Cleared Eyes. Invest Ophthalmol Vis Sci. 2018;59: 2371–2380.

14. Watson AM, Rose AH, Gibson GA, Gardner CL, Sun C, Reed DS, et al. Ribbon scanning confocal for high-speed high-resolution volume imaging of brain. PLoS One. 2017;12: e0180486.

15. Akagi T, Uji A, Huang AS, Weinreb RN, Yamada T, Miyata M, et al. Conjunctival and Intrascleral Vasculatures Assessed Using Anterior-Segment Optical Coherence Tomography Angiography in Normal Eyes. Am J Ophthalmol. 2018. doi:10.1016/j.ajo.2018.08.009

16. Loewen RT, Brown EN, Scott G, Parikh H, Schuman JS, Loewen NA. Quantification of Focal Outflow Enhancement Using Differential Canalograms. Invest Ophthalmol Vis Sci. 2016;57: 2831–2838.

17. Loewen RT, Brown EN, Roy P, Schuman JS, Sigal IA, Loewen NA. Regionally Discrete Aqueous Humor Outflow Quantification Using Fluorescein Canalograms. PLoS One. 2016;11: e0151754.

18. Dang Y, Waxman S, Wang C, Parikh HA, Bussel II, Loewen RT, et al. Rapid learning curve assessment in an ex vivo training system for microincisional glaucoma surgery. Sci Rep. 2017;7: 1605.

19. Parikh HA, Loewen RT, Roy P, Schuman JS, Lathrop KL, Loewen NA. Differential Canalograms Detect Outflow Changes from Trabecular Micro-Bypass Stents and Ab Interno Trabeculectomy. Sci Rep. 2016;6: 34705.

20. Fryirs KA, Brierley GJ. What’s in a name? A naming convention for geomorphic river types using the River Styles Framework. PLoS One. 2018;13: e0201909.

21. Katz D. Intrascleral nerve loops. Indian J Ophthalmol. 1971;19: 2–6.

22. Stevenson TC. Intrascleral Nerve Loops: A clinical study of frequency and treatment. Am J Ophthalmol. 1963;55: 935–939.

23. Reese AB. Intrascleral Nerve Loops. Trans Am Ophthalmol Soc. 1931;29: 148–153.

24. Johnstone M. 3. Intraocular pressure control through linked trabecular meshwork and collector channel motion. In: Knepper PA, Samples JR, editors. Glaucoma Research and Clinical Advances 2016 to 2018. Kugler Publications; 2016. p. 41.

25. Rosenquist R, Epstein D, Melamed S, Johnson M, Grant WM. Outflow resistance of enucleated human eyes at two different perfusion pressures and different extents of trabeculotomy. Curr Eye Res. 1989;8: 1233–1240.

26. Goldstein IJ, Hughes RC, Monsigny M, Osawa T, Sharon N. What should be called a lectin? Nature. 1980;285: 66–66.

27. Liener I. The Lectins: Properties, Functions, and Applications in Biology and Medicine. Elsevier; 1986.

28. Kataoka H, Ushiyama A, Kawakami H, Akimoto Y, Matsubara S, Iijima T. Fluorescent imaging of endothelial glycocalyx layer with wheat germ agglutinin using intravital microscopy. Microsc Res Tech. 2016;79: 31–37.

29. Scruggs AK, Cioffi EA, Cioffi DL, King JAC, Bauer NN. Lectin-Based Characterization of Vascular Cell Microparticle Glycocalyx. PLoS One. 2015;10: e0135533.

30. Shah SR, Esni F, Jakub A, Paredes J, Lath N, Malek M, et al. Embryonic mouse blood flow and oxygen correlate with early pancreatic differentiation. Dev Biol. 2011;349: 342–349.

31. Sienkiewicz AE, Rosenberg BN, Edwards G, Carreon TA, Bhattacharya SK. Aberrant glycosylation in the human trabecular meshwork. Proteomics Clin Appl. 2014;8: 130–142.

32. Bentley MD, Hann CR, Fautsch MP. Anatomical Variation of Human Collector Channel Orifices. Invest Ophthalmol Vis Sci. 2016;57: 1153–1159.

33. Hann CR, Vercnocke AJ, Bentley MD, Jorgensen SM, Fautsch MP. Anatomic Changes in Schlemm’s Canal and Collector Channels in Normal and Primary Open-Angle Glaucoma Eyes Using Low and High Perfusion PressuresDistal Outflow Pathway at Low and High Pressure. Invest Ophthalmol Vis Sci. 2014;55: 5834–5841.

34. Selbach JM, Gottanka J, Wittmann M, Lütjen-Drecoll E. Efferent and afferent innervation of primate trabecular meshwork and scleral spur. Invest Ophthalmol Vis Sci. 2000;41: 2184–2191.

35. Selbach JM, Rohen JW, Steuhl K-P, Lütjen-Drecoll E. Angioarchitecture and innervation of the primate anterior episclera. Curr Eye Res. 2005;30: 337–344.

36. McDougal DH, Gamlin PD. Autonomic control of the eye. Compr Physiol. 2015;5: 439–473.

